# Application of a novel force-field to manipulate the relationship between pelvis motion and step width in human walking

**DOI:** 10.1101/636787

**Authors:** Lauren N. Heitkamp, Katy H. Stimpson, Jesse C. Dean

**Affiliations:** Department of Health Professions, Medical University of South Carolina, Charleston, SC, 29425 USA; Ralph H. Johnson VAMC, Charleston, SC, 29401, USA

**Keywords:** Biomechanics, Legged locomotion, Rehabilitation robotics

## Abstract

Motion of the pelvis throughout a step predicts step width during human walking. This behavior is often considered an important component of ensuring bipedal stability, but can be disrupted in populations with neurological injuries. The purpose of this study was to determine whether a novel force-field that exerts mediolateral forces on the legs can manipulate the relationship between pelvis motion and step width, providing proof-of-concept for a future clinical intervention. We designed a force-field able to: 1) minimize the delivered mediolateral forces (Transparent mode); 2) apply mediolateral forces to assist the leg toward mechanically-appropriate step widths (Assistive mode); and 3) apply mediolateral forces to perturb the leg away from mechanically-appropriate step widths (Perturbing mode). Neurologically-intact participants were randomly assigned to either the Assistive group (n=12) or Perturbing group (n=12), and performed a series of walking trials in which they interfaced with the force-field. We quantified the step-by-step relationship between mediolateral pelvis displacement and step width using partial correlations. Walking in the Transparent force-field had a minimal effect on this relationship. However, force-field assistance directly strengthened the relationship between pelvis displacement and step width, whereas force-field perturbations weakened this relationship. Both assistance and perturbations were followed by short-lived effects during a wash-out period, in which the relationship between pelvis displacement and step width differed from the baseline value. The present results demonstrate that the link between pelvis motion and step width can be manipulated through mechanical means, which may be useful for retraining gait balance in clinical populations.

## I. INTRODUCTION

DURING human walking, mediolateral motion of the pelvis throughout a step is predictably related to step-by-step fluctuations in step width, a phenomenon recently reviewed by Bruijn and van Dieen [1]. Essentially, step width increases with greater mediolateral displacements or velocities of the pelvis away from the stance foot during a step [2]. The relative contributions of these two factors (pelvis displacement and velocity) to step width fluctuations have been quantified using partial correlations, revealing a substantially larger contribution of displacement [3]. This relationship between pelvis motion and step width is likely due in part to passive dynamics, with the trunk and stance leg acting as an inverted pendulum under the influence of gravitational forces. However, active control also appears to play a role beyond passive geometric effects; pelvis motion predicts hip abductor activity during swing, which in turn predicts mediolateral foot placement [4].

Whereas a predictable relationship between pelvis motion and step width is consistently observed in control participants, this relationship can be weakened following a neurological injury. Among chronic stroke survivors, the partial correlation between pelvis displacement and step width (ρ_disp_) is lower for steps taken with the paretic leg than steps taken with the non-paretic leg [5], an indirect indication that this relationship is not purely due to passive dynamics. Indeed, the contribution of active control appears to be disrupted in stroke survivors with clinically-identified balance deficits, as the link between pelvis motion and hip abductor activity during swing is weakened [6].

Varying step width to account for pelvis motion has long been proposed as an important strategy for ensuring bipedal walking balance [7]. Based on the presumed importance of this relationship, prior research has sought to manipulate pelvis dynamics and measure the resultant changes in step width. The most common approach has been to stabilize the pelvis using stiffness or damping, which directly reduces mediolateral motion of the pelvis and causes participants to prefer to walk with narrower steps [8-13]. More recently, electromechanical devices have been used to perturb or amplify the mediolateral motion of the pelvis during walking. Discrete mediolateral perturbations produce consistent shifts in foot placement and step width in the direction of the perturbation [14-15], such as a wider step when the pelvis is perturbed toward the swing leg. However, the response to extended periods of perturbation or amplified motion appears to be more complex, with reported increases [16], decreases [17], and no change [13] in step width. Despite this interest in the link between pelvis motion and step width in various mechanical contexts, no methods currently exist to produce controlled manipulations of step width itself – as will likely be necessary to determine whether this gait parameter is truly under active, dynamics-dependent control. To address this gap, we have developed a novel elastic force-field able to apply mediolateral forces to the swing leg based on pelvis motion during walking [18].

In the present study, the control of our force-field is directly based on an empirically quantified relationship between mediolateral pelvis motion during a step and the subsequent step width in neurologically-intact controls [3]. Essentially, we calculate a “mechanically-appropriate location” for each step width, based on the preceding pelvis motion (e.g. a relatively wide step if the pelvis is far medially from the stance foot; a relatively narrow step if the pelvis is near the stance foot). In some cases, we then apply mediolateral forces to the swing leg to *assist* users in achieving this step width. In other cases, we apply mediolateral forces to the swing leg to *perturb* users away from this step width (e.g. pushing the leg to take a narrow step when a wide step is mechanically appropriate).

The purpose of this study was to determine whether a novel elastic force-field can manipulate the relationship between pelvis motion and step width among neurologically-intact adults. In the longer term, our goal is to use our device to shape this relationship in clinical populations with balance deficits. Here, we hypothesized that applying forces to assist step width toward a mechanically-appropriate location would strengthen this relationship (quantified with ρ_disp_), whereas applying forces to perturb step width away from a mechanically-appropriate location would weaken this relationship. While not hypothesis-driven, our analyses also explored whether the effects of our force-field varied over time, as well as whether the effects persisted once the assistance or perturbations ceased. As part of this early-stage project, we compared the effects of several candidate equations used to identify a mechanically-appropriate step width in each step, as detailed in the Methods.

## II. METHODS

### A. Force-field Design and Control

We used a custom force-field (Fig. 1a-b) to influence step width while participants walked on a treadmill. The setup used to create this force-field has been described in detail previously [18], and consists of two steel wires in series with extension springs, running parallel to the treadmill belts. These wires interface with the legs of participants by passing through cuffs worn on the lateral sides of the shanks, with the cuffs designed to allow free anteroposterior and vertical motion of the legs relative to the wires. The wire endpoints are anchored to linear actuators (UltraMotion; Cutchoge, NY, USA) that can be rapidly repositioned mediolaterally. The net effect of this design is an elastic force-field, with mediolateral forces acting on the leg that are linearly proportional to the mediolateral distance between the leg cuff and the actuator position. For the present study, the effective mediolateral stiffness was 180 N/m.

**Fig. 1.**
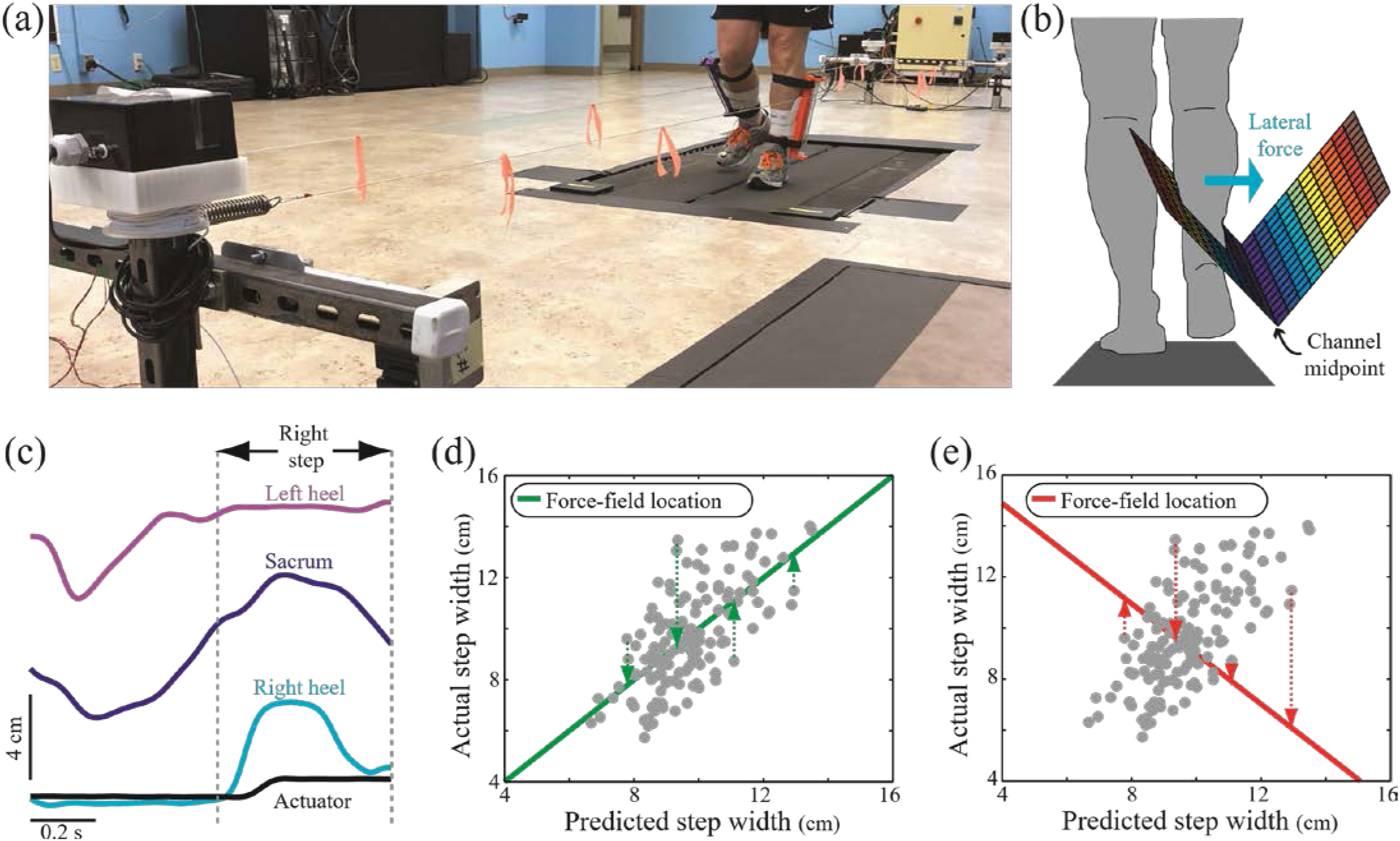
Our force-field was designed to encourage step widths that can be varied on a step-by-step basis. (a) A participant is shown walking while interfaced with the force-field. The anterior-side frame of our device is seen in the left foreground of the image. (b) The force-field takes the form of a V-shaped “channel” running parallel to the treadmill belts, schematically illustrated as a 3D surface in a frontal plane view. Actuator location controls the channel midpoint, and thus the mediolateral leg forces experienced by users during swing. (c) Mediolateral locations of the bilateral heels, sacrum, and right side actuator are illustrated for a single stride in one participant. The actuator location can be quickly repositioned during the course of a step (here, moving medially during a right step), allowing control over the mediolateral forces. (d) At the start of each step, the upcoming mechanically-appropriate step width can be predicted from a combination of mediolateral pelvis displacement and velocity (Predicted step width; x axis). In Assistive mode, the force-field moves to be aligned with this Predicted step width, thus pushing the leg’s Actual step width (y axis) toward this location. The location of the force-field (and thus the step width it encourages) is illustrated with the thick green line. Each gray dot represents a hypothetical individual step in an individual participant, comparing the predicted step widths at the start of a step with the actual step widths at the end of a step. The mediolateral forces that would act on the leg at the end of the step are depicted by the dashed green arrows for a few sample steps. (e) In Perturbing mode, the force-field pushes the leg away from the predicted mechanically-appropriate step width, following a relationship with the opposite slope to the Assistive mode, as illustrated with the thick red line. Mediolateral forces acting on the leg at the end of the step are again depicted for a few steps with dashed arrows. For comparison, the same hypothetical participant steps are illustrated as in panel (d), in order to illustrate the direction of the experienced forces.

Our prior work has demonstrated that this force-field design can encourage control participants to walk with either narrower or wider steps than normally preferred [18]. In the present study, the force-field was used to influence step width on a step-by-step basis through near real-time control of the linear actuators. This control was based on the position of active LED markers (PhaseSpace; San Leandro, CA, USA) placed on the legs and pelvis of walking participants. It should be noted that while our system has a ∼110 ms delay between the marker-driven control signal and actuator position, the actuators are able to achieve a new mediolateral position (and thus apply mediolateral force on the leg) well within the duration of a typical swing phase (Fig. 1c).

Three distinct modes of force-field control were applied: Transparent; Assistive; and Perturbing. In Transparent mode, we tracked the mediolateral position of each leg cuff and kept the corresponding actuator aligned with this position, minimizing the mediolateral forces acting on the legs. This mode allowed us to test whether simply being interfaced with our device altered the walking pattern. In Assistive and Perturbing modes, we identified the start of each step in near real-time, by tracking when an LED marker (PhaseSpace; San Leandro, CA, USA) placed on the heel switched from moving posteriorly to moving anteriorly [19]. To avoid false event detection due to marker “jitter”, we required three consecutive samples (sampled at 120 Hz) of the marker moving posteriorly prior to the detection of anterior motion – a method validated during pilot testing. For the Assistive mode, we predicted a mechanically-appropriate step width at the start of each step, and pushed the swing leg toward this location (Fig. 1d). For example, if at the start of a right step the pelvis was displaced relatively far to the right of the stance foot, we would push the swing leg to the right to take a wide step. In Perturbing mode, we again predicted a mechanically-appropriate step width at the start of each step, but pushed the swing leg away from this location (Fig. 1e). Here, if at the start of a right step the pelvis was displaced relatively far to the right of the stance foot, we would push the swing leg to the left to take a narrow step.

For both Assistive and Perturbing modes, the “mechanically-appropriate step width” was calculated based on the state of the pelvis at the start of each step. Five candidate equations for this calculation were tested (Table 1), involving various combinations of sacrum mediolateral displacement (relative to the stance heel), sacrum mediolateral velocity, and the individual participant’s mean step width. The first equation (“Personalized”) was derived from each participant’s behavior during an initial treadmill trial (as detailed below). The remaining equations (#1-4) were derived from the previously determined group-average behavior of neurologically-intact control participants walking at 1.2 m/s [3], regressing each step width value against the input variables of interest. The purpose of testing these five candidate equations was to determine whether group-based equations have similar effects as Personalized equations. This is an important consideration for future applications of this device, which will seek to promote the group-based “typical” relationship between pelvis motion and step width in populations who lack this gait behavior.

**TABLE I.** FORCE-FIELD CONTROL EQUATION

| Eq. | Assistive Mode | Perturbing Mode |
| --- | --- | --- |
| P | $A \cdot x_{sacrum} + B \cdot v_{sacrum} + C \cdot SW_{mean}$ | $-A \cdot x_{sacrum} - B \cdot v_{sacrum} + (2-C) \cdot SW_{mean}$ |
| 1 | $1.12 \cdot x_{sacrum} + 65$ | $-1.12 \cdot x_{sacrum} + 211$ |
| 2 | $1.12 \cdot x_{sacrum} + 0.47 \cdot SW_{mean}$ | $-1.12 \cdot x_{sacrum} + 1.53 \cdot SW_{mean}$ |
| 3 | $1.30 \cdot x_{sacrum} + 0.18 \cdot v_{sacrum} + 81$ | $-1.30 \cdot x_{sacrum} - 0.18 \cdot v_{sacrum} + 195$ |
| 4 | $1.30 \cdot x_{sacrum} + 0.18 \cdot v_{sacrum} + 0.59 \cdot SW_{mean}$ | $-1.30 \cdot x_{sacrum} - 0.18 \cdot v_{sacrum} + 1.41 \cdot SW_{mean}$ |

Candidate equations used to predict mechanically-appropriate step widths, and either assist or perturb participants in stepping to these locations. Here, $x_{sacrum}$ is the mediolateral displacement of the sacrum from the stance heel at the start of a step, $v_{sacrum}$ is the mediolateral velocity of the sacrum at the start of a step, and $SW_{mean}$ is the participant’s mean step width. For $x_{sacrum}$ and $v_{sacrum}$ , the direction away from the stance leg is defined as positive. For the Personalized equation (equation P), the A, B, and C coefficients are calculated for each individual participant during a Normal walking trial. For equations 1-4, these coefficients were determined based on previously quantified group-average empirical relationships. All equations use units of mm or mm/s.

### B. Experimental Procedure

This study involved 24 young, neurologically-intact participants (16 F / 8 M; age = 24 ± 2 yrs; height = 172 ± 11 cm; mass = 74 ± 13 kg; mean ± s.d.). Participants were randomly assigned to either the Assistive group (n=12) or the Perturbing group (n=12). All participants provided written informed consent using a document approved by the Medical University of South Carolina Institutional Review Board, and consistent with the Declaration of Helsinki.

Participants performed a series of treadmill walking trials at 1.2 m/s, a typical preferred walking speed in neurologically-intact controls. For all trials, participants wore a harness attached to an overhead rail that did not support body weight, but would have prevented a fall in case of a loss of balance. In the first trial, participants walked for 5-minutes under normal conditions to become accustomed to walking on the treadmill. Data from the final 2-minutes of this trial were used to calculate each participant’s mean step width (*SW*_*mean*_), and to generate the parameters for the Personalized equation for each individual participant (A, B, and C coefficients in Table 1). This was done by regressing step width values against the pelvis displacement and velocity at the start of each step.

Participants then performed a series of two 5-minute trials and five 10-minute trials (corresponding to the five force-field control equations listed in Table 1), in randomized order. In one 5-minute trial, participants walked under Normal conditions, without interfacing with the force-field. In the other 5-minute trial, participants walked with the force-field in Transparent mode for the entire trial. The structure of the remaining 10-minute trials depended on whether the participant was assigned to the Assistive or Perturbing group (n=12 per group). For the Assistive group, the force-field was in Assistive mode for the first 5-minutes of each of these trials, before switching to Transparent mode for the final 5-minutes. For the Perturbing group, the force-field was in Perturbing mode for the first half of the trial, and Transparent mode for the second half. The final 5-minutes of these trials in Transparent mode was considered a wash-out period, in which minimal mediolateral forces were applied to the legs. In order to investigate the potential effects of time during these trials in which the force-field switched modes, gait data were analyzed in 1-minute blocks.

### C. Data Collection and Processing

Active LED markers were placed bilaterally over the PSIS, ASIS, heel, lateral ankle malleolus, lateral aspect of the midfoot, and second toe. Markers were also placed on the sacrum and bilaterally on the leg cuffs, aligned with the location where the force-field wire passes through the cuffs. The present analyses focus on the sacrum and heel markers. Marker location was sampled at 120 Hz, and low-pass filtered at 10 Hz.

Gait events of interest were identified using marker data. Specifically, the start of a step was defined as the time point when the ipsilateral heel marker changed from moving posteriorly to anteriorly [19], whereas the end of a step was defined as the time point when the contralateral heel marker changed from moving posteriorly to anteriorly. The sacrum marker was used to estimate mediolateral pelvis location, as our previous work found this simplification to have only a minimal effect on our calculations [3]. Throughout each step, we measured mediolateral displacement of the pelvis relative to the stance heel and mediolateral velocity of the pelvis. We measured step width as the mediolateral displacement between the ipsilateral heel marker at the end of the step and the contralateral heel marker at the start of the step.

The primary metric of interest for this study was the partial correlation between mediolateral pelvis displacement and step width (ρ_disp_), accounting for variation in mediolateral pelvis velocity. This metric was chosen based on the specific function of the force-field in the present study; the force-field control was explicitly designed to manipulate step width based on pelvis motion, so our goal was to quantify whether the relationship between pelvis motion and step width indeed changed. Our choice of this metric was further justified based on our prior findings that: 1) predictions of step width are dominated by pelvis displacement during steady-state walking, with only a secondary effect of pelvis velocity [3]; 2) chronic stroke survivors exhibit significant paretic side deficits in the relationship between pelvis displacement and step width, but not in the relationship between pelvis velocity and step width [5]. Beyond our primary metric of ρ_disp_, previous work has also quantified the relationship between pelvis motion and step width using linear regressions (R^2^ magnitude) and the partial correlation between pelvis velocity and step width (ρ_vel_). For comparison by interested readers, the effects of our force-field on these metrics are presented in Supplementary Material.

We calculated ρ_disp_ across all steps within each minute of walking, sufficient time to reach a plateau [3]. To calculate ρ_disp_ over the course of a step, we first resampled the mediolateral pelvis displacement and velocity traces during each step to create 101-sample vectors. This resampling normalized pelvis motion as a percentage of step duration, allowing comparisons across steps of variable periods. For each normalized time point in a step (from 0-100), we then calculated ρ_disp_ as the partial correlation between all the pelvis displacement values at that time point and all the step width values at the end of the step, accounting for pelvis velocity values at that time point. This analysis process created a trajectory of ρ_disp_ values over the step duration, as done previously [3, 5].

While we calculated ρ_disp_ throughout the step, our statistical analyses focused on ρ_disp_ magnitude at the start of the step, as the swing foot was leaving the ground. This choice was based on our previous finding that muscle activity early in the swing phase had a major influence on mediolateral foot placement [4], as well as the apparent importance of this relationship early in a step in order to achieve fast walking speeds [3].

### D. Statistics

To determine whether the Transparent force-field mode truly had a minimal effect on gait, we first compared ρ_disp_ during the 5-minute Normal walking trial and the 5-minute Transparent walking trial, across all 24 participants. Following conditional equivalence testing methods [20], we first used a paired t-test to test for a significant difference between these conditions. In the case of no significant difference, we then tested for equivalence of means using two one-sided tests (TOST) procedures, with an equivalence margin equal to half the standard deviation of the ρ_disp_ value during Normal walking.

To investigate the effects of the force-field in Assistive mode, we calculated the change in ρ_disp_ magnitude relative to the baseline 5-minute Transparent trial (Δ ρ_disp_). This allowed us to focus on changes in ρ_disp_ relative to baseline, rather than on differences across individual participants. To account for any potential changes across time, ρ_disp_ was calculated for each minute of walking. We performed a repeated-measures two-way ANOVA with interactions, testing for effects of time (minutes 1-5 of walking) and control equation (see Table 1) on the change in ρ_disp_. We further performed identically structured ANOVA to test for effects of time and control equation on the change in ρ_disp_ relative to baseline during the 5-minute washout period following the Assistive period (minutes 6-10), during the 5-minute Perturbing period (minutes 1-5), and during the 5-minute washout period following the Perturbing period (minutes 6-10). In the case of a significant effect (p<0.05) we performed Tukey-Kramer post hoc tests to identify specific differences between 1-minute periods or control equations, as appropriate. All results are illustrated to display the 95% confidence interval of Δ ρ_disp_ relative to baseline. A 95% confidence interval that does not overlap with zero is interpreted as a significant difference relative to baseline.

## III. RESULTS

### A. Effects of Force-Field in Transparent Mode

Compared to Normal walking, the Transparent force-field had a minimal effect on the relationship between pelvis displacement and step width. Over the course of a step, ρ_disp_ generally increased, essentially overlapping between Normal and Transparent conditions (Fig. 2). Based on pelvis displacement at the start of each step, a paired t-test found no significant difference (p=0.30) in ρ_disp_ between Normal (ρ_disp_ = 0.68±0.07) and Transparent (ρ_disp_ = 0.67±0.07) conditions. Given an equivalence margin of 0.04 (half the standard deviation during Normal walking), a TOST analysis found significant (p=0.008) evidence for equivalence between these conditions. Across participants, the 95% confidence interval for the difference in ρ_disp_ between the Normal and Transparent conditions was [-0.01 to 0.03].

**Fig. 2.**
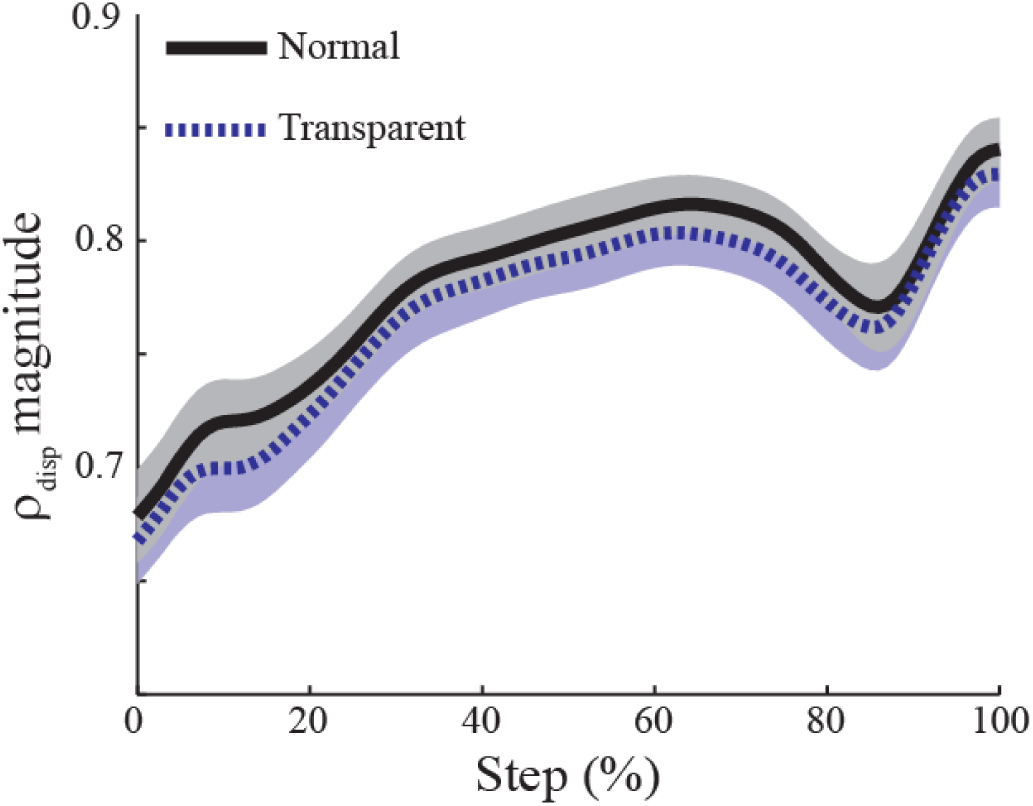
The relationship between pelvis displacement and step width was quite similar for Normal and Transparent walking conditions, as quantified using ρ_disp_ and illustrated from the start of a step (0%) to the end of a step (100%). The lines indicate the mean values of these metrics, and the shaded areas indicate 95% confidence intervals.

### B. Effects of Force-Field in Assistive Mode

Force-field assistance caused clear increases in the strength of the relationship between pelvis displacement and step width (Fig. 3a). The increases in ρ_disp_ magnitude relative to baseline were consistent across the 5-minutes of walking in Assistive mode (Fig. 3b), with no significant main effect of time (p=0.10). However, force-field control equation did have a significant main effect (p<0.0001) on the increase in ρ_disp_ magnitude, as equations that did not include pelvis velocity (i.e. Equations 1 and 2) generally produced smaller increases (Fig.3c). No significant interaction between time and control equation was observed (p=0.09).

**Fig. 3.**
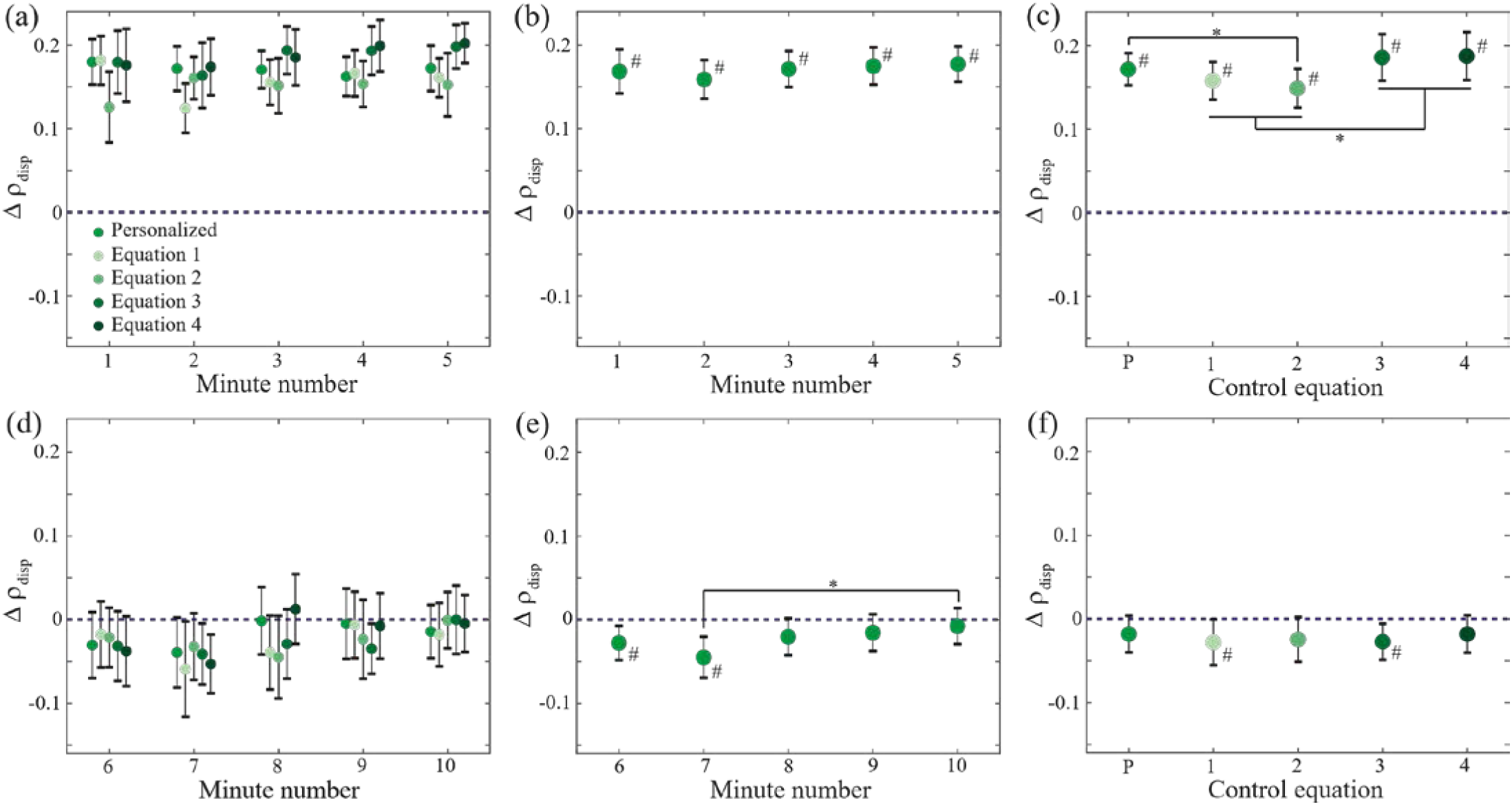
Force-field assistance influenced the relationship between pelvis displacement and step width. The top row (panels a-c) focuses on the changes in ρ_disp_ during the 5-minute period in which assistance was applied (minutes 1-5), whereas the bottom row (panels d-f) focuses on the subsequent 5-minute washout period (minutes 6-10). Panels (a) and (d) illustrate Δ ρ_disp_ for each minute of walking and each control equation. As no significant interactions between time and control equation were observed, we illustrate the statistical main effects of time in panels (b) and (e), and the main effects of control equation in panels (c) and (f). All panels illustrate the mean difference in ρ_disp_ relative to the baseline Transparent trial (here indicated by the dashed horizontal line), with error bars indicating 95% confidence intervals. Asterisks (*) indicate a significant post-hoc difference between the indicated values, and pound signs (#) indicate a significant difference from baseline.

Following cessation of force-field assistance, the relationship between pelvis displacement and step width did not immediately return to its baseline level during the subsequent wash-out period (Fig. 3d). The change in ρ_disp_ magnitude relative to baseline varied significantly (p=0.024) over time (Fig. 3e), with a value lower than baseline for the first 2-minutes of walking before returning to its original level. During this wash-out period, the change in ρ_disp_ magnitude was not significantly influenced by a main effect of force-field equation (p=0.86; Fig. 3f) or an interaction between time and control equation (p=0.84).

### C. Effects of Force-Field in Perturbing Mode

Force-field perturbations caused a general decrease in the strength of the relationship between pelvis displacement and step width (Fig. 4a), although these changes were of smaller magnitude than those observed with force-field assistance. The decreases in ρ_disp_ magnitude were not significantly affected by a main effect of time (p=0.82; Fig. 4b), a main effect of control equation (p=0.27; Fig. 4c), or an interaction between time and control equation (p=0.44).

**Fig. 4.**
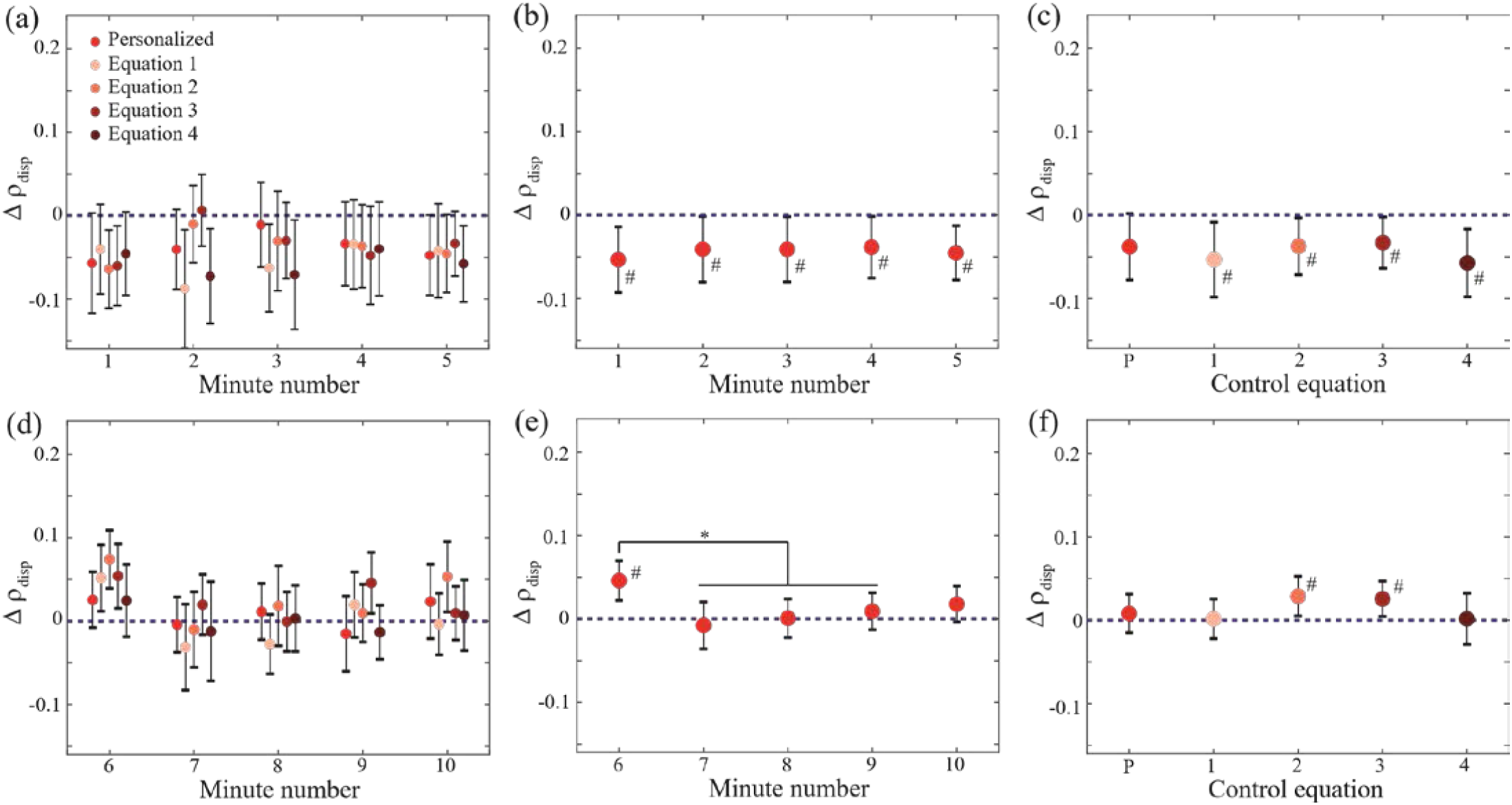
Force-field perturbations influenced the relationship between pelvis displacement and step width, generally in the opposite direction to the effect observed with force-field assistance. This figure follows the structure used in Fig. 3 above, with the top row (panels a-c) focusing on changes in ρ_disp_ during the 5-minute period in which perturbations were applied, and the bottom row (panels d-f) focusing on the subsequent 5-minute washout period.

As with force-field assistance, the relationship between displacement and step width did not immediately return to its baseline level once perturbations ceased (Fig. 4d). A significant main effect of time (p=0.0001) was observed, as ρ_disp_ magnitude was higher than its baseline level for the first minute of walking without perturbations, before returning to baseline (Fig. 4e). Control equation also had a significant main effect (p=0.039) on ρ_disp_ magnitude during the washout period, although none of the individual post-hoc comparisons between equations reached significance (Fig. 4f). No significant interaction between time and control equation was observed (p=0.47).

### D. Force-Field Effects throughout a Step

While our statistical analyses focused on the effects of our force-field on ρ_disp_ at the start of a step (as justified in the Methods), we here describe ρ_disp_ values throughout the course of a step. Force-field assistance increased ρ_disp_ magnitude throughout a step, although the magnitude of this increase relative to baseline declined as the step progressed (Fig. 5a). The negative effects on ρ_disp_ during the first 2 minutes of the washout period were only observed early in a step, with ρ_disp_ returning to its baseline level by approximately midway through a step (Fig. 5a). In contrast, the decreases in ρ_disp_ produced by force-field perturbations were of similar magnitude throughout the course of a step, possibly even increasing as the step progressed (Fig. 5b). During the first minute of the subsequent wash-out period, ρ_disp_ generally remained higher than its baseline level throughout the step (Fig. 5b). We did not observe any clear effects of force-field control equation on these patterns of ρ_disp_ changes throughout a step.

**Fig. 5.**
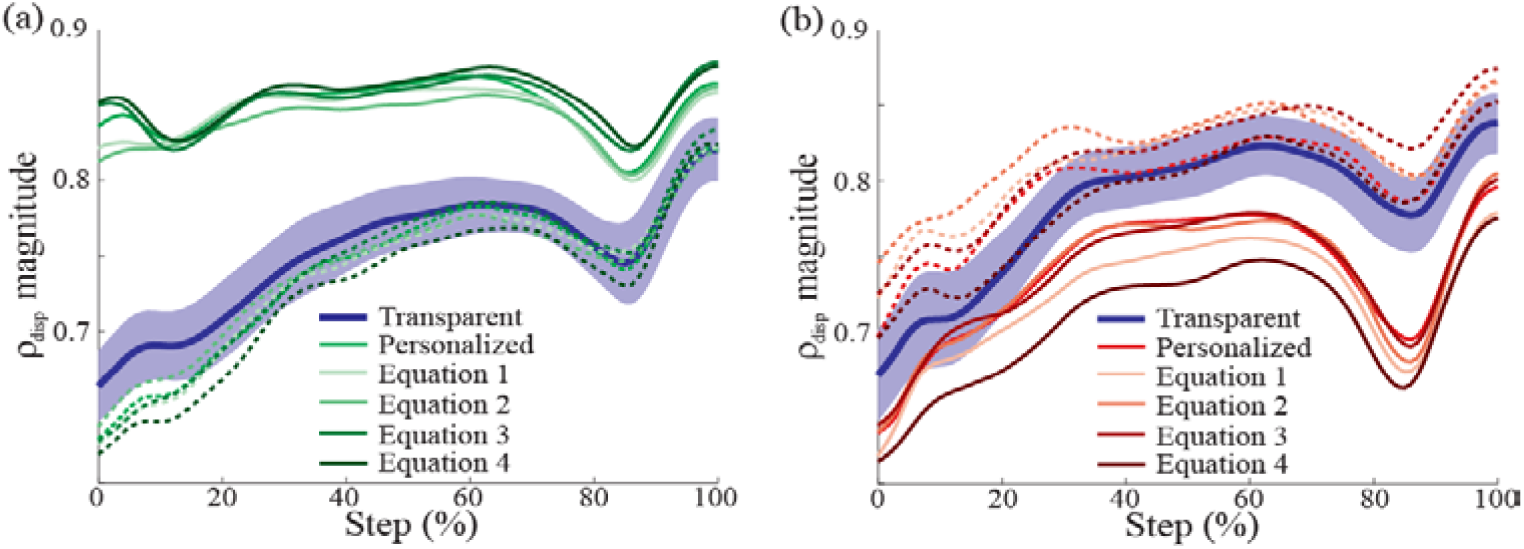
The effects of each force-field mode on ρ_disp_ are illustrated throughout a step, in comparison to the Transparent condition. Panel illustrates the effect of force-field assistance, while panel (b) illustrates the effects of force-field perturbations. For both panels, the solid lines show ρ_disp_ while in Assistive or Perturbing mode. The dashed lines show ρ_disp_ at the beginning of the subsequent washout period (first 2-minutes for Assistive mode; first 1-minute for Perturbing mode. To avoid extensive overlap, the 95% confidence interval is shown only for the Transparent condition, as the shaded blue area.

## IV. DISCUSSION

Our novel elastic force-field effectively manipulated the relationship between pelvis displacement and step width. As hypothesized, force-field assistance strengthened this relationship, whereas force-field perturbations weakened this relationship. Following cessation of the force-field assistance or perturbations, participants did not immediately return to their baseline behavior, instead exhibiting an altered relationship between pelvis displacement and step width for 1-2 minutes. The control equation used to identify a mechanically-appropriate step width location had only a minor effect on gait behavior, which was fairly consistent within each force-field mode.

In essence, our force-field did what it was designed to do – predictably either increasing or decreasing the partial correlation between pelvis displacement and step width. These effects were consistently observed across the 5-minute walking periods in which forces were applied, with no apparent decay over time. While hypothesized, the observed changes in ρ_disp_ were not guaranteed, as participants could have feasibly adjusted their active control to resist the relatively weak mediolateral forces during a step (∼15 N for even the largest perturbation illustrated in Fig. 1e), thus holding ρ_disp_ constant across conditions. However, the smaller magnitude of the changes directly evoked by perturbations (Δ ρ_disp_ = −0.04 ± 0.08; mean ± s.d. across control equations and time) compared to assistance (Δ ρ_disp_ = +0.17 ± 0.05) suggests that participants did not entirely ignore the applied forces. Although speculative, perhaps participants were more likely to adjust their active control to resist the forces produced in Perturbing mode. This type of response would be consistent with the recent finding that humans are more likely to adapt their gait pattern in response to mechanical contexts that challenge balance [21]. If participants did not adjust their control, the Perturbing force-field would encourage narrower than typical steps for large pelvis displacements (possibly increasing the risk of a lateral loss of balance [22]), and wider than necessary steps for small pelvis displacements (with associated increases in mechanical, muscular, and metabolic demands [23-24]).

While not the primary focus of this study, the altered gait behavior early in wash-out periods provides further indirect evidence for an adjustment in the underlying active control. The short-lived (1-2 minutes) overshoot in ρ_disp_ relative to baseline following a change in mechanical context is qualitatively similar to after-effects commonly observed in studies of gait adaptation, such as with split-belt walking [25]. Perhaps the decrease in ρ_disp_ following force-field assistance is a sign that participants “slacked” [26] in terms of their pelvis-dependent active control of step width. Conversely, perhaps the increase in ρ_disp_ following force-field perturbations was caused by an increased contribution of active control to the relationship between pelvis displacement and step width. To provide some context, the average increase in ρ_disp_ during the first minute of walking after being perturbed was 0.05. For comparison, among chronic stroke survivors this metric was an average of 0.09 lower for steps taken with the paretic leg than steps taken with the non-paretic leg [5]. Further work is needed to determine whether force-field perturbations of the type investigated here have the potential to strengthen the link between pelvis motion and paretic step width in this clinical population, which may help to improve some aspects of walking balance. As part of this future work, the development of methods that would allow the quantification of ρ_disp_ outside of the laboratory (possibly using inertial measurement units) could have clear benefits for extending this research to clinical settings.

While intriguing, the potential indicators of adaptation observed during wash-out periods should be interpreted cautiously due to limitations of our experimental and analytical design. For example, repeated exposure to a novel mechanical environment can influence the presence and magnitude of after-effects in later bouts [27]. The present study involved multiple bouts of force-field Assistance or Perturbations, but with the underlying control equation presented in randomized order to reduce potential ordering effects. This structure prevents a simple direct comparison across walking bouts. Additionally, prior work has generally identified the largest adaptation-driven changes in gait behavior using only a few strides (∼5) following cessation of the altered mechanical context [28]. This analysis is not possible with the present study’s primary metric, which requires an extended period of walking for accurate quantification [3]. Despite these limitations, the present results motivate future work to more thoroughly investigate potential adaptation of the step-by-step fluctuations in step width.

Participants responded similarly to force-field equations based on their individual behavior (Personalized condition) and based on previously quantified group average behavior. This is an important finding for our long-term goal of applying our force-field methods to neurologically-injured populations who exhibit deficits in the relationship between pelvis motion and step width [5]. Our electromechanical force-field is likely too complex to be implemented in most clinical settings, but can serve as a “device emulator” to identify the force-field characteristics and level of complexity needed to evoke the targeted step-by-step changes in step width – similar to a recently developed prosthesis emulator designed to speed the development of novel prostheses and exoskeletons [29]. The present results suggest that the forces acting on the legs need not account for an individual’s exact gait parameters or mediolateral pelvis velocity in order to effectively manipulate the relationship between pelvis displacement and step width. This result suggests that a simpler (perhaps entirely passive) device that exerts mediolateral forces on the swing leg based on the mediolateral location of the pelvis could be an effective training tool.

Overall, the present study demonstrated that a novel elastic force-field can predictably alter the relationship between pelvis displacement and step width. This relationship is generally believed to be important for walking stability [1] and is disrupted in a patient population with impaired balance [5], but is not directly targeted by existing gait and balance training strategies for clinical populations. The present work can thus serve as a foundation for future clinical applications, in which training of mechanically-appropriate adjustments in step width may be added to current clinical paradigms. In the long-term, repeated exposure to an altered mechanical environment may have the potential to produce retained changes in the link between pelvis motion and step width, paralleling previously observed improvements in post-stroke step length asymmetry following repeated exposure to split-belt walking [30-31].

## ACKNOWLEDGEMENTS

The authors would like to thank Abby Kelly for statistical assistance.

