## Supplementary Material for "Application of a novel force-field to manipulate the relationship between pelvis motion and step width in human walking"

The results presented in this manuscript's main text describe the effects of our novel force-field on $\rho_{disp}$, the partial correlation between mediolateral pelvis displacement at the start of a step and step width, while accounting for mediolateral pelvis velocity at the start of a step. Our primary focus on this metric was based on prior research showing that pelvis displacement is more strongly predictive of step width than pelvis velocity [1], and that chronic stroke survivors exhibit clear deficits only in the relationship between pelvis displacement and step width [2]. However, other metrics have been used to quantify the relationship between pelvis motion and step width. Here, we present the results of secondary analyses focused on two other previously reported metrics [1-2].

### I. Regression-Based $R^2$

For each participant and each minute of walking for each condition, we performed a linear regression to quantify the extent to which pelvis motion predicted the eventual step width. Specifically, mediolateral pelvis displacement and velocity at the start of each step were included as independent variables, and step width was included as the dependent variable. We quantified the ability of the pelvis metrics to predict step width using the magnitude of the $R^2$ from this regression. Our statistical analyses investigating the effects of time and force-field control equation are identical to those performed using $\rho_{disp}$ and described in the manuscript main text.

#### A. *Effects of Force-Field in Assistive Mode*

Overall, the effects of force-field assistance on $R^2$ magnitude were highly similar to those reported in the main text for $\rho_{disp}$. The Assistive mode clearly increased $R^2$ magnitude (Fig. S1a). The evoked increases in $R^2$ magnitude were fairly consistent across the 5-minute period (Fig. S1b), with no significant main effect of time (p=0.11). As with $\rho_{disp}$, the force-field control

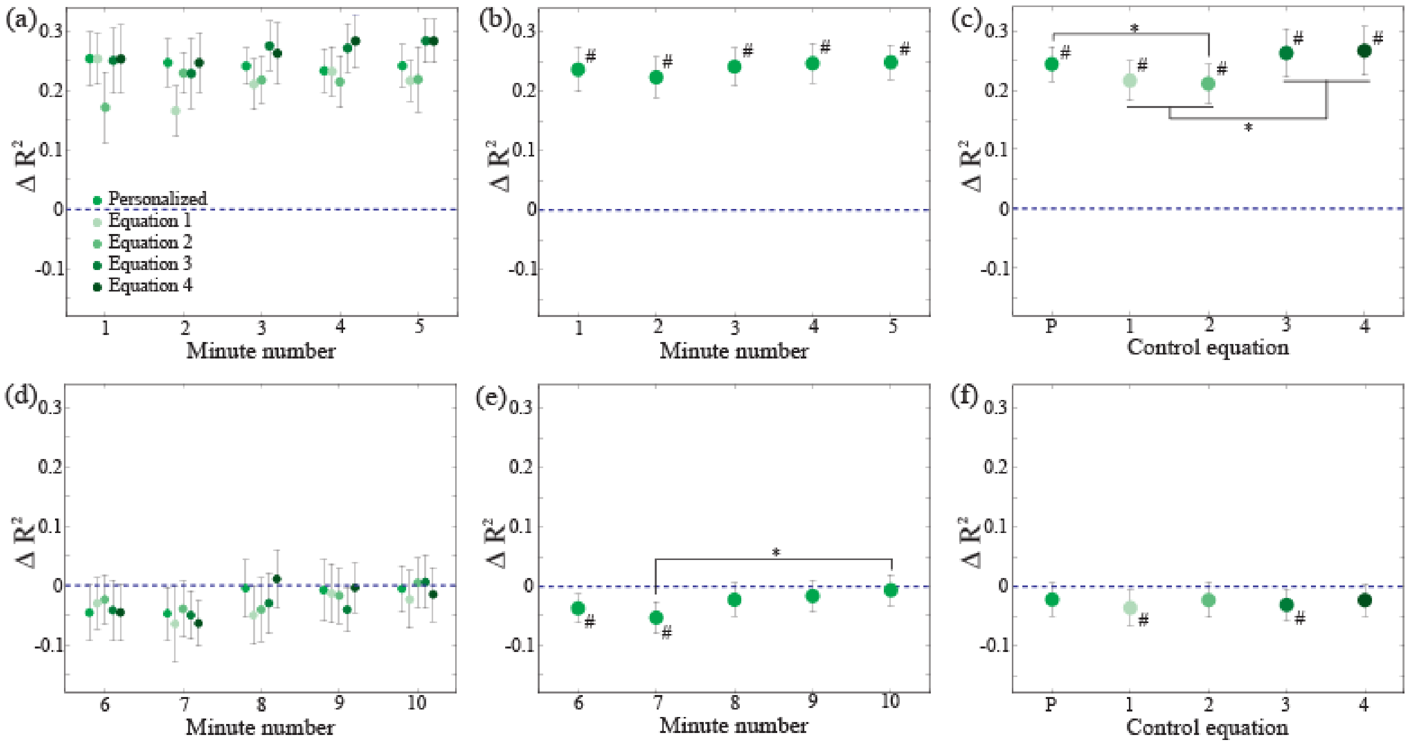


Fig. S1. Force-field assistance influenced $R^2$ magnitude, both during and after its application. The top row (panels a-c) focuses on the changes in $R^2$ ($\Delta R^2$) during the 5-minute period in which assistance was applied (minutes 1-5), whereas the bottom row (panels d-f) focuses on the subsequent 5-minute washout period (minutes 6-10). Panels (a) and (d) illustrate $\Delta R^2$ for each minute of walking and each control equation. The statistical main effects of time are illustrated in panels (b) and (e), and the main effects of control equation are illustrated in panels (c) and (f). All panels illustrate the mean difference in $R^2$ relative to the baseline Transparent trial (here indicated by the dashed horizontal line), with error bars indicating 95% confidence intervals. Asterisks (*) indicate a significant post-hoc difference between the indicated values, and pound signs (#) indicate a significant difference from baseline.

equation had a significant main effect ($p<0.0001$) on the increase in $R^2$, as equations that included pelvis velocity (P, 3, and 4) produced larger increases (Fig. S1c). A significant interaction between time and control equation was present ($p=0.040$), as Equations 1 and 2 produced only relatively small increases in $R^2$ magnitude during minutes 2 and 1, respectively.

The changes in $R^2$ magnitude after the force-field assistance ceased also directly paralleled those seen for $\rho_{disp}$ magnitude (Fig. S1d). $R^2$ magnitude did not immediately return to its baseline level after the assistance stopped, but instead varied significantly over time ($p=0.007$), remaining reduced in comparison to its baseline level for 2-minutes (Fig. S1e). During the wash-out period, $R^2$ magnitude was not significantly affected by the control equation ($p=0.80$; Fig. S1f) or an interaction between time and control equation ($p=0.91$)

*B. Effects of Force-Field in Perturbing Mode*

The effects of force-field perturbations on $R^2$ magnitude were again highly similar to those seen for $\rho_{disp}$. Such perturbations generally decreased $R^2$ magnitude in comparison to the baseline level (Fig. S2a). These decreases were not significantly affected by time ($p=0.42$), remaining fairly consistent across the 5-minute walking period (Fig. S2b). The observed effects on $R^2$ were also not significantly affected by the force-field control equation ($p=0.11$), with similar magnitude decreases observed for all equations (Fig. S2c). No significant interaction between time and control equation was observed ($p=0.50$).

During the washout period after perturbations ceased, the pattern of changes in $R^2$ magnitude was nearly identical to that reported for $\rho_{disp}$ (Fig. S2d). A significant main effect of time was present ($p=0.0005$), as $R^2$ magnitude was significantly higher than its baseline level for the first minute of walking without perturbations (Fig. S2e). Force-field control equation also had a significant main effect ($p=0.046$) on the change in $R^2$ magnitude during the washout period, although none of the individual post-hoc comparisons reached significance (Fig. S2f). No significant interaction between time and control equation was present ($p=0.33$).

### II. Pelvis Velocity Partial Correlation

For each participant and each minute of walking for each condition, we calculated the partial correlation between the mediolateral pelvis velocity at the start of the step and the eventual step width ($\rho_{vel}$), accounting for the mediolateral pelvis displacement at the start of each step. This calculation thus mimicked the calculation of our primary outcome variable ($\rho_{disp}$), but with a focus on pelvis velocity rather than displacement.

*A. Effects of Force-Field in Assistive Mode*

The effects of force-field assistance on $\rho_{vel}$ magnitude were highly variable (Fig. S3a), producing notable increases for only a subset of cases. While the force-field was in Assistive mode, time had a significant main effect ($p=0.0004$) on the change in $\rho_{vel}$ relative to baseline, with greater increases later in the 5-minute walking period (Fig. S3b). The change in $\rho_{vel}$ was also significantly affected ($p<0.0001$) by a main effect of control equation. Perhaps unsurprisingly, only control equations that included pelvis velocity (equations P, 3, and 4) produced significant increases in $\rho_{vel}$ (Fig. S3c). No significant interaction between time and control equation was observed ($p=0.41$).

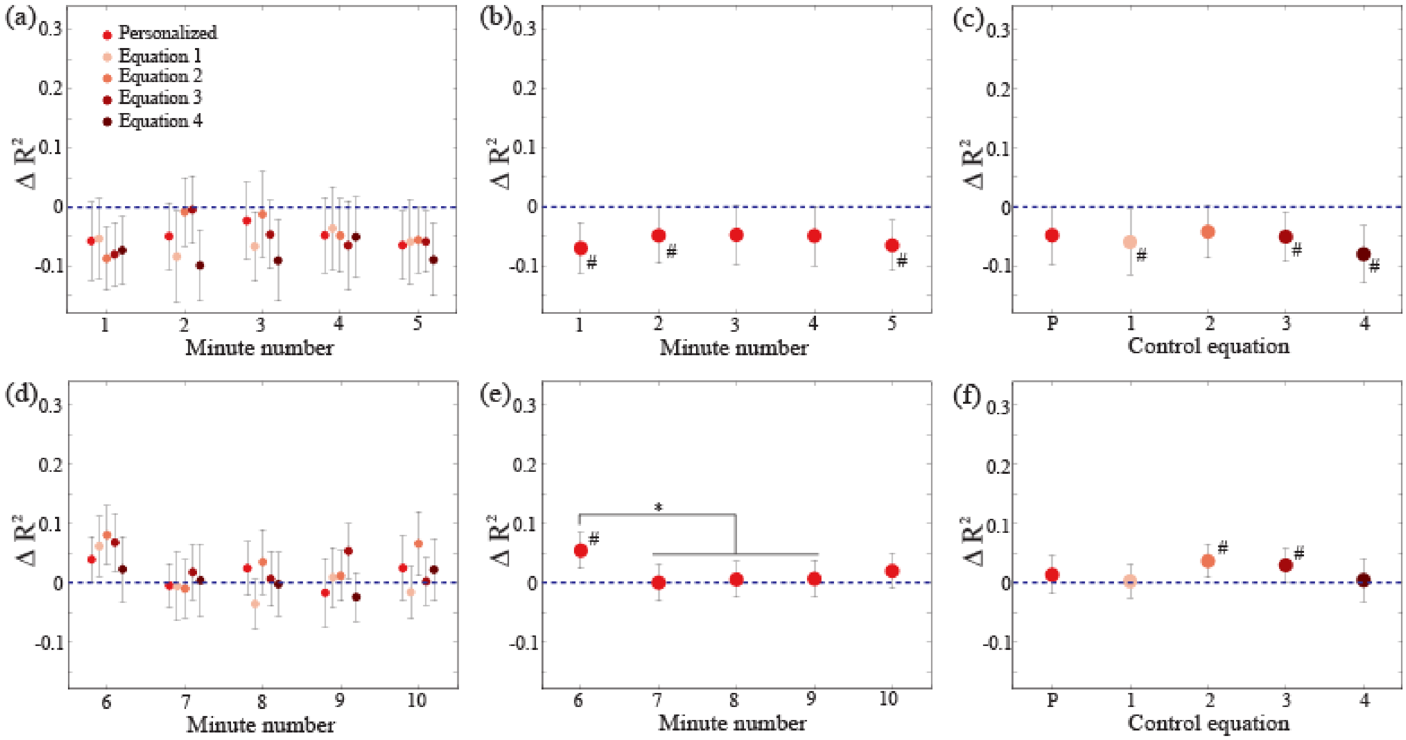


Fig. S2. Force-field perturbations influenced $R^2$ magnitude, both during and after their application. The structure of this figure parallels that of Figure S1.

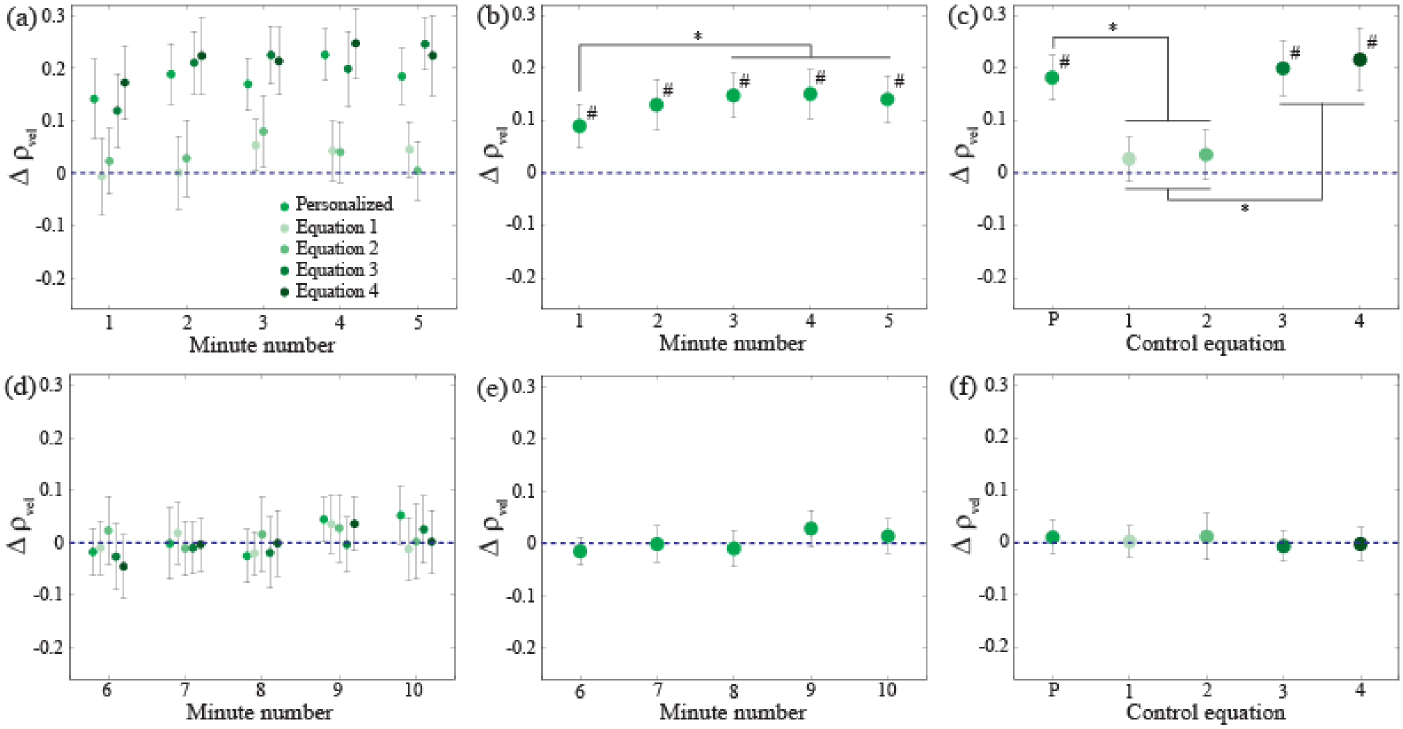


Fig. S3. Assistance only influenced $\rho_{vel}$ during its application, and for control equations that included pelvis velocity. The figure structure parallels Figure S1.

Unlike the previously presented metrics of $\rho_{disp}$ and $R^2$, $\rho_{vel}$ did differ from its baseline value during the washout period (Fig. S3d). Across this 5-minute period, $\rho_{vel}$ was not significantly affected by a main effect of time ($p=0.07$; Fig. S3e), control equation ($p=0.79$; Fig. S3f), or an interaction between time and control equation ($p=0.90$)

### *B. Effects of Force-Field in Perturbing Mode*

The effects of force-field perturbations on $\rho_{vel}$ magnitude were also quite variable (Fig. S4a), only producing clear decreases in this metric for a subset of cases. While perturbations were applied, the decrease in $\rho_{vel}$ was significantly affected by time ($p=0.031$), as $\rho_{vel}$ generally increased after the first minute of walking (Fig. S4b). The

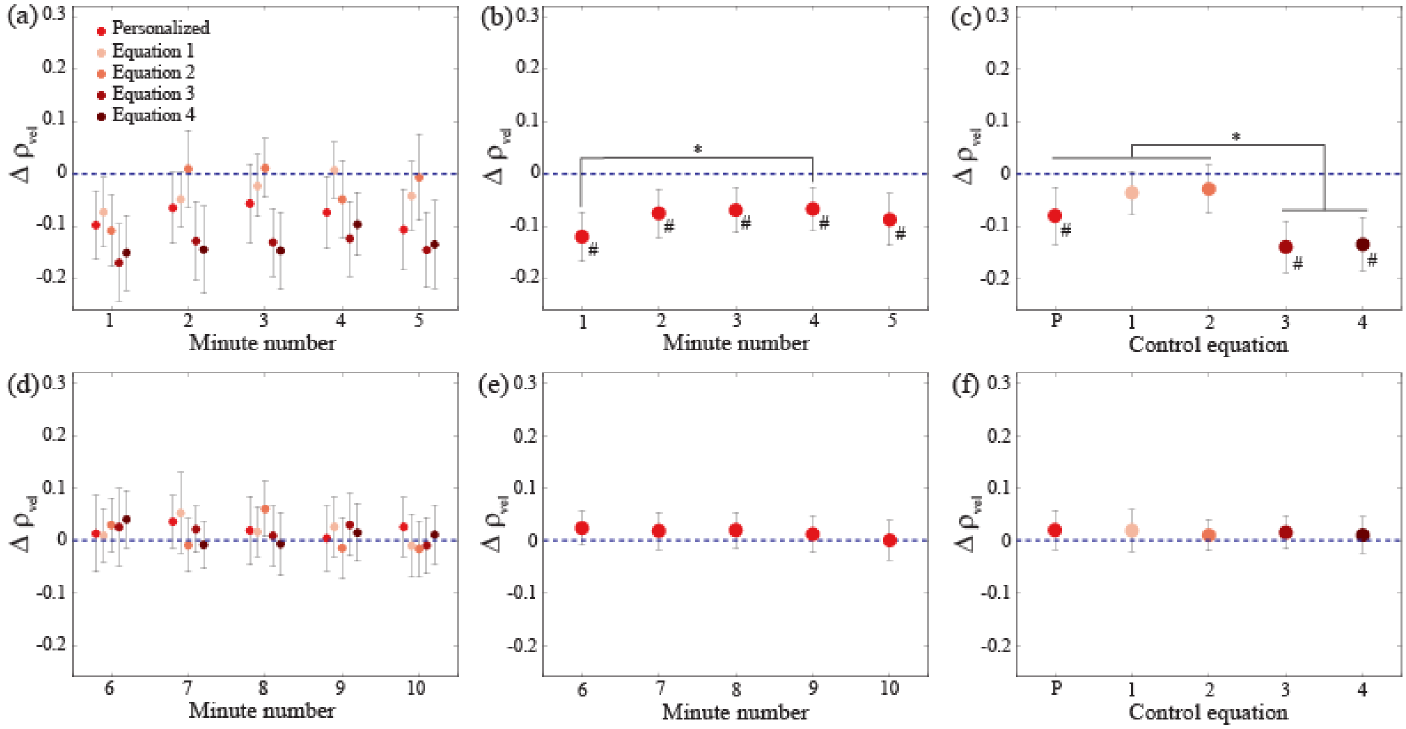


Fig. S4. Perturbations only influenced $\rho_{vel}$ while they were applied, and for control equations that included pelvis velocity. The figure structure parallels Figure S1.

changes in $\rho_{vel}$ relative to baseline were also significantly affected by control equation ($p<0.0001$). Significant decreases in $\rho_{vel}$ were only produced by control equations that included velocity (P, 3, and 4) (Fig. S4c). No significant interaction between time and control equation was present ($p=0.81$).

As with force-field assistance, force-field perturbations did not produce noticeable changes in $\rho_{vel}$ during the subsequent washout period (Fig. S4d). During this period, $\rho_{vel}$ was not significantly affected by a main effect of time ($p=0.68$; Fig. S4e), control equation ($p=0.97$; Fig. S4f), or an interaction between time and control equation ($p=0.79$).

### III. Summary

The effects of our novel force-field on the step-by-step relationship between pelvis motion and step width can be quantified using several metrics. The effects revealed using the partial correlation between mediolateral pelvis displacement and step width ($\rho_{disp}$) closely parallel those seen with the $R^2$ magnitude calculated using a linear regression involving both pelvis displacement and velocity. These similarities were present both while assistance or perturbations are applied and during the subsequent washout period. The similarity between these metrics is likely attributable to variation in pelvis displacement dominating the subsequent variation in step width, as has been previously reported [1]. In contrast, the force-field's effects on the partial correlation between pelvis velocity and step width ($\rho_{vel}$) differed substantially from the other quantified metrics. This metric appears to only be affected by control equations that directly include pelvis velocity, and did not exhibit after-effects during the washout period once the force-field assistance or perturbations ceased.
